# Multi-batch cytometry data integration for optimal immunophenotyping

**DOI:** 10.1101/2020.07.14.202432

**Authors:** Masato Ogishi, Rui Yang, Conor Gruber, Simon Pelham, András N. Spaan, Jérémie Rosain, Marwa Chbihi, Ji Eun Han, V Koneti Rao, Leena Kainulainen, Jacinta Bustamante, Bertrand Boisson, Dusan Bogunovic, Stéphanie Boisson-Dupuis, Jean-Laurent Casanova

## Abstract

We describe the integration of multi-batch cytometry datasets (iMUBAC), a flexible, robust, and scalable computational framework for unsupervised cell-type identification across multiple batches of high-dimensional cytometry datasets. After overlaying cells from healthy controls across multiple batches, iMUBAC learns batch-specific cell-type classification boundaries and identifies aberrant immunophenotypes in patient samples. We illustrate unbiased and streamlined immunophenotyping, using both in-house and public mass and flow cytometry datasets.

## Main

High-dimensional cytometry — including mass cytometry (*i.e*., cytometry by time-of-flight, CyTOF) and spectral flow cytometry (*e.g*., Cytek Aurora) — facilitates the phenotyping of precious samples from patients for various immune cell subsets, at single-cell resolution, which is of particular interest in human immunology. Ideally, a large number of samples should be processed simultaneously to ensure comparability. However, this is not always possible. For example, patients may be recruited across the globe over decades and longitudinally monitored in prospectively expanding cohorts for rare diseases, such as inborn errors of immunity^1^. Similarly, some investigations may involve multiple pilot studies with small numbers of patients followed by larger-scale validation studies. In such situations, the integration of multiple batches of experiments processed on different occasions and at different sites is inevitable. The simplest solution for multi-batch integration is to gate cell subsets with manual batch-to-batch adjustments. However, manual analyses of cytometry data are inherently subjective, knowledge-driven, and non-scalable for multiple batches of datasets. More objective, unbiased, and scalable methods are therefore desired.

Efforts have been made to facilitate high-dimensional data inspection and unsupervised cell-subset identification (*e.g*., viSNE^2^, SPADE^3^, FlowSOM^4,5^, CITRUS^6^, and CellCNN^7^). However, these automated approaches are themselves sensitive to the batch effects resulting from the separate processing of different experiments. Per-channel (single-dimensional) signal intensity normalization has been attempted (*e.g*., cydar^8^, CytoNorm^9^, and CyTOFBatchAdjust^10^) as a means of overcoming batch effects. However, single-dimensional normalization cannot fully account for cell type-specific high-dimensional batch effects. Moreover, batch correction on the cells of patients with aberrant immunophenotypes is undesirable, due to inherent uncertainty about over-or under-correction. The requirement for identical technical replicates across all batches poses another challenge in CytoNorm and CyTOFBatchAdjust^9,10^. A more high-dimensional approach, SAUCIE^11^, was recently described. SAUCIE automates batch correction and clustering, but its clustering resolution is inadequate for rare subsets. For example, its analyses of T cells from subjects with acute dengue virus infection and healthy controls do not identify a distinct cluster corresponding to CD4^+^CD25^+^FOXP3^+^ regulatory T cells (Tregs). Thus, current computational approaches do not accomplish high-dimensional batch correction and unbiased immunophenotyping with sufficient resolution from multiple batches of datasets.

Here, we present iMUBAC (integration of multi-batch cytometry datasets), a flexible, robust, and scalable computational framework for rational interbatch comparisons through high-dimensional batch correction and unsupervised cell-type identification across multiple batches (Fig. 1). The workflow can be broken down into four steps. First, iMUBAC performs a data-adaptive, automated preprocessing, such as excluding doublets and dead cells and ensuring inter-batch consistency in panel design. Second, iMUBAC batch-corrects cells from healthy local controls, but not from travel/family controls or patients, using Harmony^12^ to reduce batch effects before clustering. No technical replicates are required. Third, iMUBAC performs unsupervised clustering with the batch-corrected expression values, with i) FlowSOM^4^ followed by metaclustering with ConsensusClusterPlus or ii) dimension reduction by uniform manifold approximation and projection (UMAP)^13^ followed by shared nearest-neighbor (SNN) graph-based clustering^14^. If desired, the clusters can be further manually merged and identified to improve interpretability in subsequent analyses. Fourth, iMUBAC trains batch-specific classifiers through machine learning with non-corrected expression values. Here, the idea is to “back-propagate” cell-type annotations defined in the batch-corrected high-dimensional space into the non-corrected, batch-specific spaces in which the patients’ cells are embedded. The entire workflow is independent of patients (and of travel/family controls), for which immunophenotypes need to be determined, thereby circumventing potential over- or under-correction.

**Figure 1.**
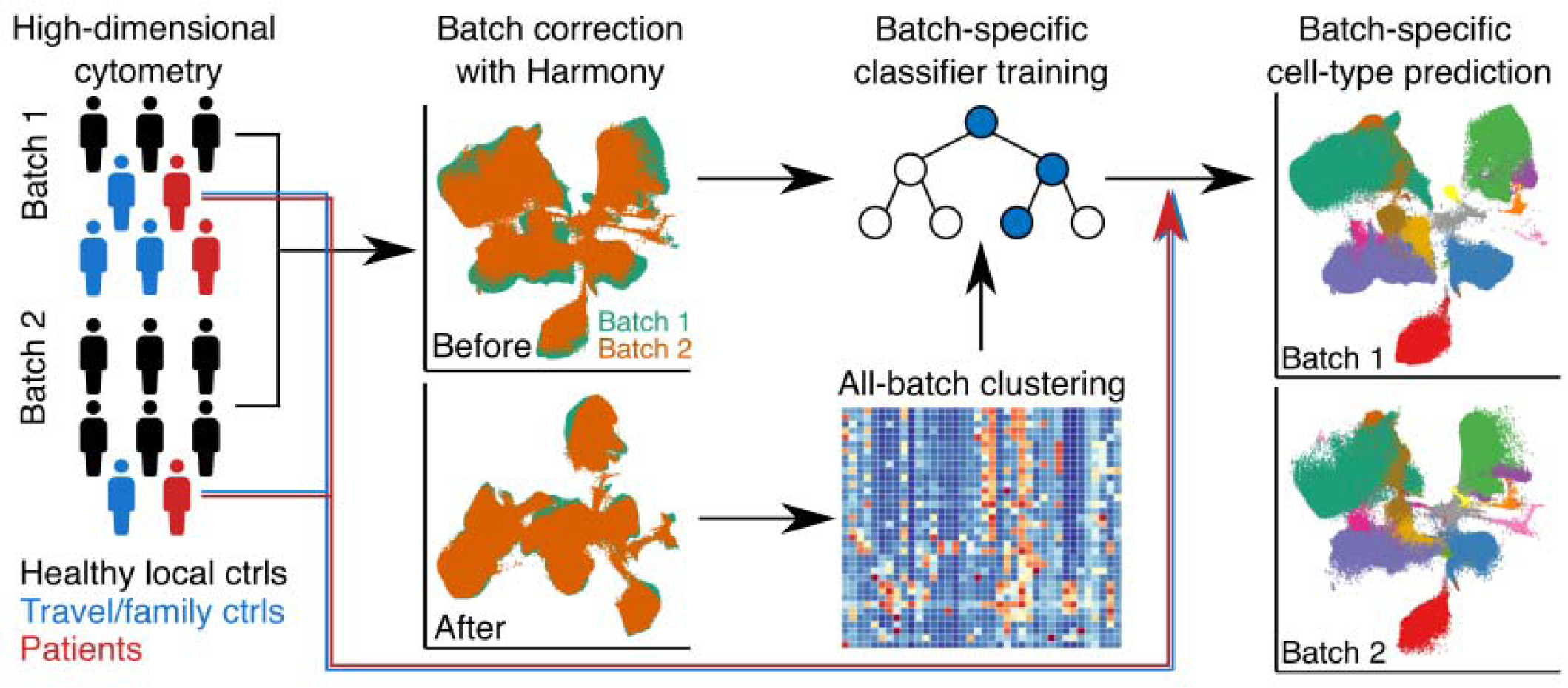
Integration of multi-batch cytometry datasets (iMUBAC). Multiple cytometry datasets can be integrated for rational inter-batch and inter-individual comparisons. Even experiments with heterogeneous designs (*e.g*., different in the numbers of local controls and patients) or inconsistent panels without shared technical replicates can be integrated. Only cells from healthy local controls were used for batch correction. The batch-corrected expression values were then used for unsupervised clustering, followed by manual annotation if desired. The non-batch-corrected expression values, tied to cell-type annotations, were then used to train batch-specific cell-type classifiers. Finally, cell types were predicted for the rest of the cells, including the cells of patients and their travel/family controls.

We first tested iMUBAC on in-house multi-batch CyTOF datasets for peripheral blood mononuclear cells (PBMCs) from patients with unusually severe infectious diseases and their travel/family controls (75 individuals, 6 batches, 38 surface markers). After batch-correction with healthy local controls as “anchors,” FlowSOM/ConsensusClusterPlus defined 60 metaclusters, which we merged and identified manually (Fig. S1-3). We calculated the percentages of the various subsets among live single leukocytes (CD45^-^CD66b^+^ cells, excluding granulocytes). The cell-type frequencies of both technical (*i.e*., experiments performed on different dates with aliquots of identical biological materials, *N*=2) and biological (*i.e*., experiments performed on different dates with biological materials obtained from identical donors on different occasions, *N*=2) duplicates correlated well between the two batches tested (Fig. S4). Moreover, the estimated and manually determined frequencies of both abundant (*e.g*., αβ T cells) and rare (*e.g*., innate lymphoid cells) cell subsets correlated reasonably well across a wide range of percentages (from ∼0.01% to ∼70%), not only among local controls but also among travel/family controls and patients (Fig. S5). Collectively, iMUBAC rationally integrates multi-batch cytometry datasets, to identify cell types consistent with state-of-the-art gating in both controls and patients.

We then applied iMUBAC to in-house multi-batch spectral flow cytometry datasets for PBMCs (14 individuals, 2 batches, 14 cell-surface and 4 intracellular markers). The datasets included data from patients with two monogenic forms of autoimmunity: FAS deficiency^15,16^ and *STAT3* gain-of-function^17–19^, both of which are known to be accompanied by lymphoproliferation and high counts of circulating CD4^-^CD8^-^ double-negative αβ T (DN T) cells. After manually gating out dead cells and doublets, we defined 50 metaclusters with FlowSOM/ConsensusClusterPlus, which we merged and identified manually (Fig. 2a, 2b, and S6). As expected, we observed an expansion of a cluster representing DN T cells in patients with FAS deficiency and, to a lesser extent, in those with *STAT3* gain-of-function (Fig. 2c). Conventional flow cytometry with a different antibody panel validated this expansion of DN T cells (Fig. S7). iMUBAC also revealed a decrease in the levels of Vδ2 γδ T, CD8^+^ mucosal-associated invariant T (MAIT), and CD16^+^ natural killer (NK) cells in both groups, possibly reflecting a previously unappreciated level of immunopathological homogeneity between these two inborn errors of immunity (Fig. 2c). iMUBAC can be applied to spectral flow cytometry datasets, as well as CyTOF datasets, and readily identifies both known and unappreciated immunophenotypes in patients with rare immunological disorders.

**Figure 2.**
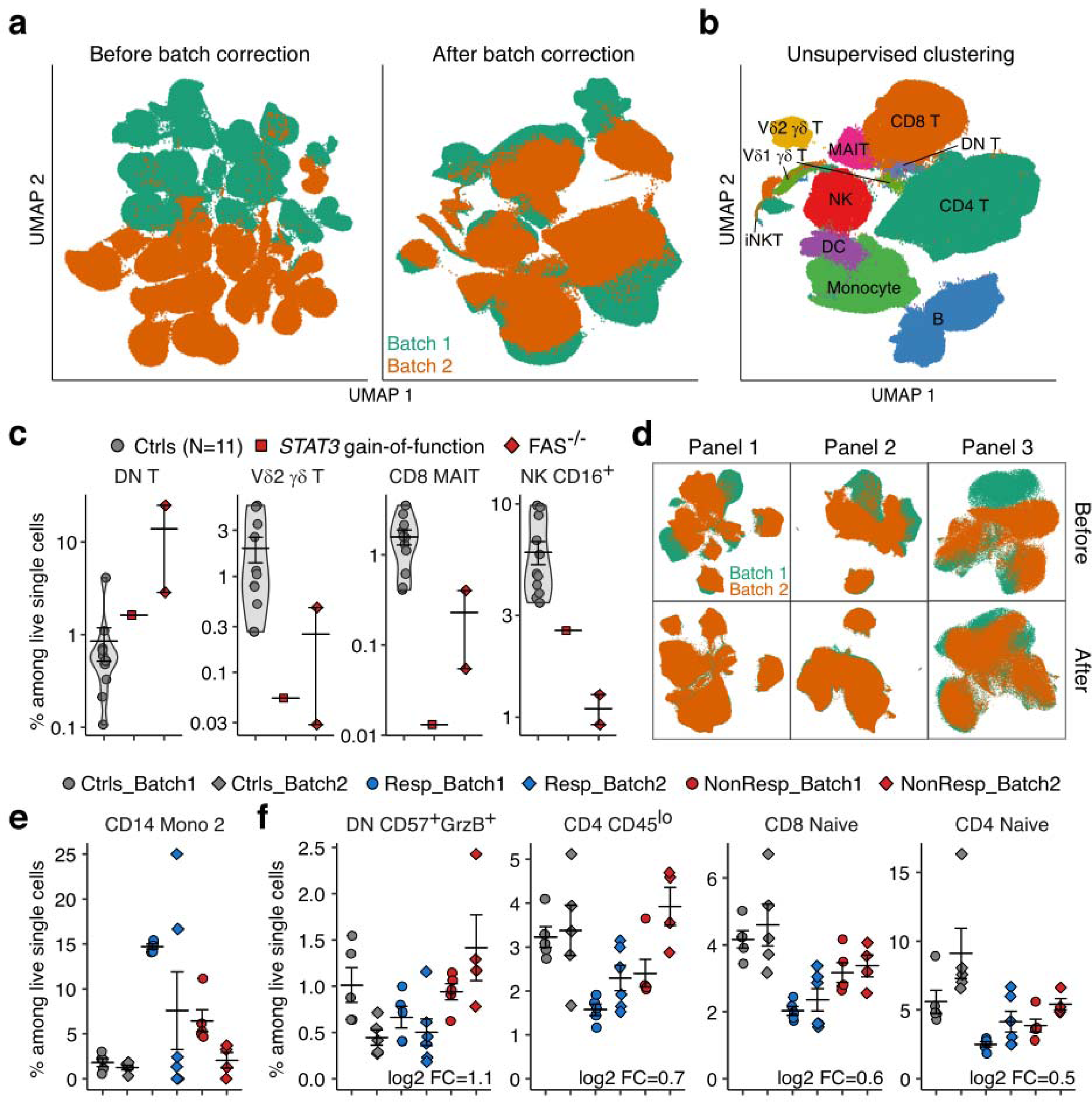
iMUBAC readily identifies both known and unknown immunophenotypes. (a-c) Spectral flow cytometry datasets for peripheral blood mononuclear cells (PBMCs) from patients with monogenic forms of autoimmunity. (a) Batch correction. (b) Cell-type identification through unsupervised clustering. (c) Selected immunophenotypes of patients with FAS deficiency and *STAT3* gain-of-function mutations. (d-f) CyTOF datasets for PBMCs from patients on PD-1 blockade treatments^20^. The original study compared immunophenotypes between responders and non-responders and reported higher levels of CD14^+^CD16^-^HLA-DR^hi^ monocytes among responders at baseline. (d) Batch correction. Panel 1, T-cell phenotyping. Panel 2, cytokines. Panel 3, myeloid phenotyping. (e and f) Differential abundance (DA) analysis. All subsets from the three panels were simultaneously tested for DA with *edgeR*^21^ after adjustment for treatment and batch effects. Percentages among live single leukocytes before PD-1 blockade are shown. (e) Expansion of the “CD14 Mono 2” cluster, corresponding to CD14^+^CD16^-^HLA-DR^hi^CD86^hi^PD-L1^hi^ monocytes, in responders. (f) DA for subsets identified statistically significant when only pretreatment datasets were used for DA testing. Estimated log2 fold-changes between responders and non-responders are also shown.

Finally, we applied iMUBAC to previously published CyTOF datasets for PBMCs from 10 healthy controls and 20 patients with stage IV melanoma before and after PD-1 blockade immunotherapy (11 responders and nine non-responders, respectively) that had been stained with three antibody panels^20^. We analyzed two batches simultaneously (*i.e*., discovery and validation cohorts). Healthy controls were used for batch correction, clustering with FlowSOM/ConsensusClusterPlus, and classifier training (Fig. 2d and S8-10). We tested the differential abundance (DA) between responders and non-responders using the quasi-likelihood F-test (QLF) in *edgeR*^21^, as previously applied to CyTOF analysis^8,22^. Eight DA subsets were identified, with statistical significance after adjustment for treatment status and batch effects. In particular, the “CD14 Mono 2” cluster, corresponding to CD14^+^CD16^-^HLA-DR^hi^CD86^hi^PD-L1^hi^ monocytes, was expanded in responders consistent with the findings of the original report^20^ (Fig. 2e). Moreover, four DA subsets, all T-cell subsets, were reproducibly identified even if only pretreatment datasets were used (Fig. 2f). These subsets could potentially be used as biomarkers of a better prognosis, before the initiation of PD-1 blockade immunotherapy. iMUBAC can be used to streamline an exploratory immunophenotyping analysis in clinical pilot studies of common disease conditions.

High-dimensional cytometry has considerably improved human immunological studies, despite limited sample availability. Moreover, platforms such as FlowRepository or Cytobank facilitate the sharing of cytometry data, making it possible to foster discoveries through meta-analysis. We anticipate that this flexible, robust, and scalable workflow, available on GitHub (https://github.com/casanova-lab/iMUBAC), will expedite rational comparative analyses of multi-batch cytometry datasets and facilitate novel discoveries through unbiased and streamlined immunophenotyping.

## Methods

### Cells

Peripheral blood mononuclear cells (PBMCs) were isolated from heparinized venous blood samples by Ficoll-Hypaque density gradient centrifugation (GE Healthcare). Cells were cryopreserved in bovine fetal calf serum supplemented with 10% dimethyl sulfoxide (DMSO) and stored at -150°C until use. Patients with various severe infectious diseases were included in this study. When blood samples from patients were collected at distant sites, the blood samples were transported to either the Paris or the New York branch of our laboratory overnight and processed. We accounted for the effect of blood sample transportation, by collecting samples from healthy volunteers or healthy family members and transporting and processing them simultaneously with the transported samples (travel/family controls). Cells from healthy local volunteers collected and processed locally were also used (local controls). We included multiple local controls in each cytometry experiment.

### Spectral flow cytometry

Two experiments were performed on separate dates. We studied frozen PBMCs from 11 locally recruited healthy adult controls and three patients (two patients with homozygous loss-of-function mutations of *FAS* and one patient with a heterozygous gain-of-function *STAT3* mutation). Freshly thawed PBMCs (2×10^6^ cells for controls and 5×10^6^ cells for the patient) were stained with the Zombie NIR Fixable Viability Kit (BioLegend, 1:2000) for 15 minutes on ice. Cells were then stained with the following panel on ice for 30 minutes: FcR Blocking Reagent (Miltenyi Biotec, 1:50), anti-CD3-BD Horizon V450 (BD Biosciences, 1:450), anti-CD4-BUV563 (BD Biosciences, 1:450), anti-CD8-BUV737 (BD Biosciences, 1:150), anti-CD14-BUV395 (BD Biosciences, 1:100), anti-CD16-PE-Dazzle (BioLegend, 1:150), anti-CD20-BV785 (BD Biosciences, 1:150), anti-CD56-BV605 (BioLegend, 1:50), anti-γδTCR-Alexa Fluor 647 (BioLegend, 1:50), anti-Vδ1-FITC (Miltenyi Biotec, 1:450), anti-Vδ2-APC-Fire750 (BioLegend, 1:1350), anti-Vα7.2-Alexa Fluor 700 (BioLegend, 1:50), MR1-BV421 (provided by the NIH Tetramer Core Facility, 1:200), anti-Vα24-Jα18-BV480 (BD Biosciences, 1:50), and anti-Vβ11-APC (Miltenyi Biotec, 1:150) antibodies. The cells were fixed on ice for 30 minutes and permeabilized with the True-Nuclear Transcription Factor Buffer Set (BioLegend). The cells were then stained for intracellular transcription factors overnight in the dark at 4°C with the following panel: anti-T-bet-PE-Cy7 (BioLegend, 1:1350), anti-GATA3-BV711 (BioLegend, 1:50), anti-RORγT-PE (BD Biosciences, 1:50), and anti-EOMES-PerCP-eFluor 710 (eBioscience, 1:50) antibodies. Cells were acquired with an Aurora cytometer (Cytek). Data were inspected using FlowJo, and manually gated live singlets were then exported as FCS files for subsequent iMUBAC analysis.

### Conventional flow cytometry

PBMCs (3×10^6^ cells) from two locally recruited healthy adult controls and three patients (two patients with homozygous null mutations of *FAS* and one patient with a homozygous gain-of-function *STAT3* mutation) were stained with the following panel on ice for 20 minutes: FcR Blocking Reagent (Miltenyi Biotec, 1:50), anti-αβTCR-PE/Cy7 (BioLegend, 1:100), anti-CD3-BV421 (BioLegend, 1:100), anti-CD4-redFluor 710 (Tonbo Biosciences, 1:100), anti-CD8-PerCP/Cy7 (BioLegend, 1:100), anti-CD14-APC/Cy7 (BioLegend, 1:100), anti-CD16-APC (BioLegend, 1:100), anti-CD19-Super Bright 645 (eBioscience, 1:100), and anti-CD56-Alexa Fluor 488 (BD Biosciences, 1:100). Cells were then stained with 7-AAD (Tonbo Biosciences, 1:200) on ice for 10 minutes. Cells were acquired with a BD FACS Aria (BD Biosciences). Compensation was performed using single-stained PBMCs as controls. Data were analyzed using FlowJo.

### Mass cytometry (CyTOF)

PBMCs from 39 healthy individuals (30 adults and nine children or adolescents), 24 patients with various severe infectious diseases and autoimmune diseases, and 12 travel/family controls were studied in six different batches of experiments. Freshly thawed PBMCs (1.0×10^6^ cells per panel) were washed with barcode permeabilization buffer (Fluidigm) and barcoded with Fluidigm’s Cell-ID 20-Plex Pd Barcoding Kit. Samples were then washed and pooled into a single tube, Fc-blocked with FcX (BioLegend) and heparin-blocked to prevent non-specific binding. Cells were then stained with a panel of metal-conjugated antibodies obtained from Fluidigm: anti-CD45-89Y, anti-CD57-113In, anti-CD11c-115In, anti-CD33-141Pr, anti-CD19-142Nd, anti-CD45RA-143Nd, anti-CD141-144Nd, anti-CD4-145Nd, anti-CD8-146Nd, anti-CD20-147Sm, anti-CD16-148Nd, anti-CD127-149Sm, anti-CD1c-150Nd, anti-CD123-151Eu, anti-CD66b-152Sm, anti-PD-1-153Eu, anti-CD86-154Sm, anti-CD27-155Gd, anti-CCR5-156Gd, anti-CD117-158Gd, anti-CD24-159Tb, anti-CD14-160Gd, anti-CD56-161Dy, anti-γδTCR-162Dy, anti-DRTH2-163Dy, anti-CLEC12A-164Dy, anti-CCR6-165Ho, anti-CD25-166Er, anti-CCR7-167Er, anti-CD3-168Er, anti-CX3CR1-169Tm, anti-CD38-170Er, anti-CD161-171Yb, anti-CD209-172Yb, anti-CXCR3-173Yb, anti-HLA-DR-174Yb, anti-CCR4-176Yb, and anti-CD11b-209Bi. After surface staining, the samples were fixed and stored until acquired on a Helios mass cytometer (Fluidigm). Dead cells and doublets were excluded by staining the cells, before and after fixation, with a rhodium-based dead cell exclusion intercalator (Rh103) and cationic iridium nucleic acid intercalators (Ir191 and Ir193), respectively.

### Public CyTOF datasets

CyTOF datasets for PBMCs from patients on PD-1 blockade^20^ were downloaded from FlowRepository (https://flowrepository.org/experiments/1124). The study from which these data were taken analyzed PBMCs from patients with melanoma before and about 12 weeks after treatment with either nivolumab or pembrolizumab. Patients were classified as responders or non-responders based on treatment outcomes for the first 15 weeks of treatment. The datasets also contained data for PBMCs from healthy donors at two corresponding time points. The datasets consisted of two batches (*i.e*., experiments processed at two separate dates) used as discovery and validation cohorts in the original study. The first batch contained data for five healthy donors, five responders, and five non-responders, whereas the second batch contained data for five healthy donors, six responders, and four non-responders. In this study, these two batches were analyzed simultaneously in an exploratory analysis.

### Computational analysis

All computational analyses were performed with R version 4.0.0 (https://www.r-project.org/)^23^ The R package *iMUBAC* and example codes are available from GitHub (https://github.com/casanova-lab/iMUBAC). Full scripts are available upon request.

### Integration of multi-batch cytometry datasets (iMUBAC)

The iMUBAC workflow consists of four steps: i) preprocessing, ii) batch correction, iii) unsupervised clustering and cell-type annotation, and iv) batch-specific cell-type prediction.

#### Preprocessing

Batch-specific preprocessing was performed as follows. First, CyTOF data files in the FCS format were imported into R with the *ncdfFlow* package. The *ncdfFlow* package can be used for the memory-efficient HDF5-based storage of cytometry data. The *truncate_max_range* option was disabled. For CyTOF data, the *transformation* option was also disabled, as the transformation implemented in the underlying *read.FCS* package is optimized for flow cytometry. Second, channel names were organized. This step resolves batch-to-batch differences in the panel design such that identical markers measured using different channels (*i.e*., fluorochrome- or metal-conjugated antibodies) are aligned. Third, for CyTOF data, doublets and dead cells were excluded in a data-adaptive manner. In this step, all cells from all samples in a single batch of an experiment were pooled, such that identical gates were applied to all samples in a given batch. For DNA-based gating, the *dnaGate* function in the *cydar* package was used, and outliers on both the higher and lower sides (considered to be doublets and debris, respectively) were excluded. For event length and dead cell exclusion dye-based gating, the *outlierGate* function in the *cydar* package was used, and outliers on the higher side were excluded. In our in-house CyTOF datasets, the intercalator Rh103 was used to exclude dead cells, whereas the intercalators Ir191 and Ir193 were used to exclude doublets and debris. In the PD-1 blockade CyTOF datasets, the intercalator Pt198 was used to exclude dead cells, whereas the intercalators Ir191 and Ir193 were used to exclude doublets and debris. For our in-house pre-gated spectral flow cytometry datasets, the channel for the Zombie NIR Fixable Viability dye was used to exclude dead cells, and automated gating for doublets and DNA content was disabled. The expression values were then transformed. For CyTOF data, a hyperbolic arcsin-transformation was applied, with a cofactor of five. For spectral flow cytometry data, Logicle transformation was applied, with parameters estimated in a data-adaptive manner with the *estimateLogicle* function implemented in the *flowCore* package. Finally, any event with zeros for all markers was discarded.

After the batch-specific preprocessing, the outputs were concatenated, with only markers common to all batches retained, to form a single *SingleCellExperiment* object in R.

#### Batch correction

The goal of this step is to enable the system to learn batch-to-batch deviations due to technical effects but not biological variabilities. We, therefore, used only data from healthy controls for batch correction, excluding data for patients and travel/family controls. Data were first down-sampled to 200,000 cells per batch (unless otherwise stated) by taking approximately equal numbers of cells from each control, to reduce the computational burden. For the in-house spectral flow datasets, we used 500,000 cells per batch, to ensure the robust identification of invariant natural killer T (iNKT) cells, an extremely rare innate-like T-cell subset. For the PD-1 blockade CyTOF Panel 3 (Myeloid Panel) dataset, we used 50,000 cells per batch, due to the low total cell counts in the dataset. We then batch-corrected expression values for all markers with Harmony^12^, using the default parameters. Rather than performing principal component analysis (PCA), we used each marker directly as an input for batch correction. The effect of batch correction was assessed by manual inspection, with uniform manifold approximation and projection (UMAP) used for data visualization^13^.

#### Unsupervised clustering and cell-type annotation

The goal of this step was to identify cell types in an unsupervised manner. We implemented two methods: i) FlowSOM-guided clustering followed by ConsensusClusterPlus-guided metaclustering and ii) UMAP-based dimension reduction followed by shared nearest neighbor (SNN) graph-based clustering^14^, as described below.

##### The FlowSOM/ConsensusClusterPlus method

This approach was inspired by the workflow described by Nowicka et al.^5^ Briefly, batch-corrected expression values were subjected to unsupervised clustering with FlowSOM^4^, using the *FlowSOM* package, followed by metaclustering with the *ConsensusClusterPlus* package. Euclidean distance was used for metaclustering. For the in-house spectral flow cytometry and CyTOF datasets, we generated 50 and 60 metaclusters, respectively, to improve the resolution of cell-type identification. For the PD-1 blockade CyTOF datasets, we generated 40 metaclusters. In addition to this clustering, we generated a heatmap for the median expression levels of all markers for each of the clusters, to facilitate the manual determination of cell identity.

##### SNN graph method

This approach was inspired by the workflow for single-cell RNA sequencing datasets implemented in the *scran* package. Batch-corrected expression values were first dimension-reduced via UMAP into ten dimensions. These dimensions were then used to construct an SNN graph with the *buildSNNGraph* function in *scran*, using the default settings. Finally, the graph was divided into communities, or clusters, with the Louvain algorithm, implemented as the *cluster_louvain* function in the *igraph* package^24^.

#### Batch-specific cell-type prediction

The goal of this step is to allow the system to learn, automatically, the cell-type classification rules, or cell-type boundaries, in a batch-specific manner, and to propagate the boundaries to all the cells in a given batch, including cells from travel/family controls and patients. Importantly, we used non-batch-corrected expression values tied to cluster labels defined in the unsupervised clustering section. First, cells from the healthy controls used for unsupervised clustering were further downsampled, retaining a maximum of 100 cells per cluster from a given batch. This step reduces the computational burden and also alleviates the class imbalance problem during machine learning, as there are both highly abundant cell subsets (*e.g*., CD14^+^ monocytes) and rare cell subsets (*e.g*., plasmacytoid dendritic cells) among human PBMCs. We tested several conditions and found that cell-type classification rules can be learned successfully from 100 events per cell-type. A classifier was then trained with the *caret* package^25^. We selected the extremely randomized trees^26^ algorithm implemented in the *extraTrees* package. After centering and scaling the non-batch-corrected expression values, we performed five-times repeated 10-fold cross-validations with internal upsampling to maximize the Kappa statistic. Hyperparameters were tuned for each batch; *mtry* was tuned from five to 15, whereas *numRandomCuts* was tuned from one to two, the ranges being empirically determined. The trained batch-specific classifier was then applied to predict clusters for all cells in a given batch, including the cells of patients and travel/family controls. This approach assumes that all cells of patients and travel/family controls fall into one of the clusters defined from the cells of healthy controls.

### Differential abundance analysis

For the PD-1 blockade CyTOF datasets, the raw counts of cell subsets identified from the three panels were tested simultaneously for differential abundance (DA) with the quasi-likelihood F-test (QLF) framework of the package *edgeR*^21^. When both pre- and post-treatment datasets were used, the DA between groups (*i.e*., responders and non-responders) was assessed with adjustment for both treatment and batch effects. When only pretreatment datasets were used, the DA between groups was assessed with adjustment for batch effects only. The DA values for subsets with an absolute log2 fold-change of at least 0.5 and an adjusted *P*-value below 0.05 were considered statistically significant.

## Acknowledgments

We would like to thank the patients, their relatives, and their physicians for participating in this study; Dominick Papandrea, Yelena Nemirovskaya, Mark Woollett, Lazaro Lorenzo-Diaz, and Cécile Patissier for administrative assistance; Tatiana Kochetkov for technical assistance; the members of the laboratory for helpful discussions. We thank the Flow Cytometry Resource Center at the Rockefeller University for technical support for flow cytometry. We also thank the Human Immune Monitoring Core at the Icahn School of Medicine at Mount Sinai for technical assistance with mass cytometry. We thank the National Institutes of Health (NIH) Tetramer Core Facility (NTCF) for providing the MR1 tetramer, which was developed jointly with Dr. James McCluskey, Dr. Jamie Rossjohn, and Dr. David Fairlie.

## Funding

M.O. was supported by the Rockefeller University PhD program, the Funai Foundation for Information Technology (FFIT), and the New York Hideyo Noguchi Memorial Society, Inc. (HNMS). R.Y. was supported by the Immune Deficiency Foundation and Stony Wold-Herbert Fund. A.N.S. was supported by the European Commission (EC, Horizon 2020 Marie Skłodowska-Curie Individual Fellowship #789645), the Dutch Research Council (NWO, Rubicon Grant #019.171LW.015), and the European Molecular Biology Organization (EMBO, Long-Term Fellowship #ALTF 84-2017, non-stipendiary). J.R. was supported by INSERM PhD program (*poste d’accueil* INSERM). The Laboratory of Human Genetics of Infectious Diseases is supported by the National Institute of Allergy and Infectious Diseases grant number 5R37AI095983, the St. Giles Foundation, the Rockefeller University, Howard Hughes Medical Institute, Institut National de la Santé et de la Recherche Médicale (INSERM), Université de Paris, the French Foundation for Medical Research (FRM) (EQU201903007798), the French National Research Agency under the “Investments for the Future” program (grant number ANR-10-IAHU-01), the Integrative Biology of Emerging Infectious Diseases Laboratory of Excellence (ANR-10-LABX-62-IBEID), the SCOR Corporate Foundation for Science, and the GENMSMD project (grant number ANR-16-CE17-0005-01 to J.B.), and SRC2017 (to J.B.).

---

**Figure S1.**
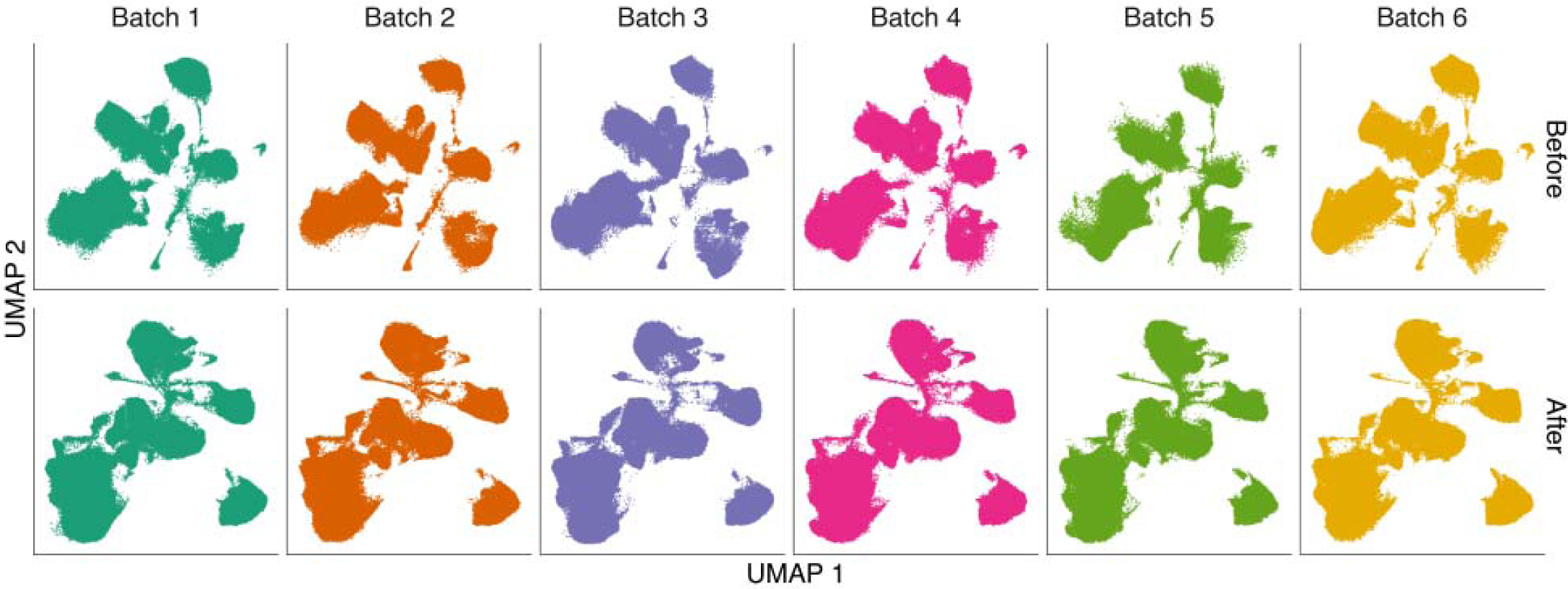
Batch correction of the in-house multi-batch CyTOF datasets. PBMCs from healthy local controls, travel/family controls, and patients with unusually severe infectious diseases (*N*=75 in total) were stained with a general immunophenotyping panel (38 surface markers) and analyzed on six different occasions. The six batches of datasets were integrated through iMUBAC. In total, 200,000 cells per batch randomly selected from healthy local controls were batch-corrected with Harmony^12^. Uniform manifold approximation and projection (UMAP)^13^ visualizations are shown.

**Figure S2.**
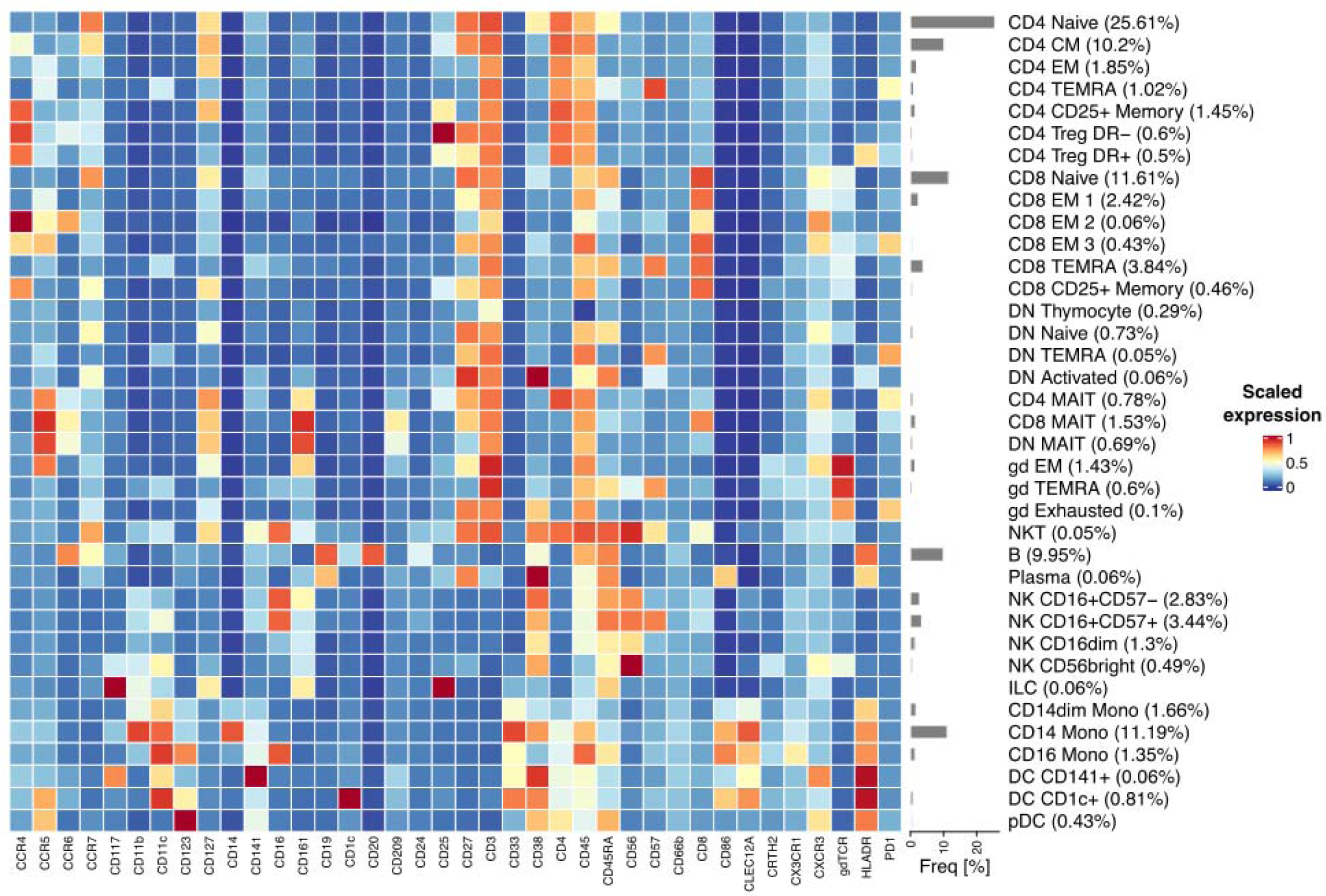
Cell-type identification for the in-house CyTOF datasets. After batch correction, the batch-corrected expression values were used for unsupervised clustering with FlowSOM^4^, followed by metaclustering with ConsensusClusterPlus. Initially, 60 metaclusters were generated. Clusters were manually inspected to determine their identity. A summary heatmap of the scaled median expression levels of all markers for all cell subsets is shown. Clusters resembling granulocytes (CD45-CD66b), eosinophils, basophils, and mast cells were excluded from the heatmap analysis.

**Figure S3.**
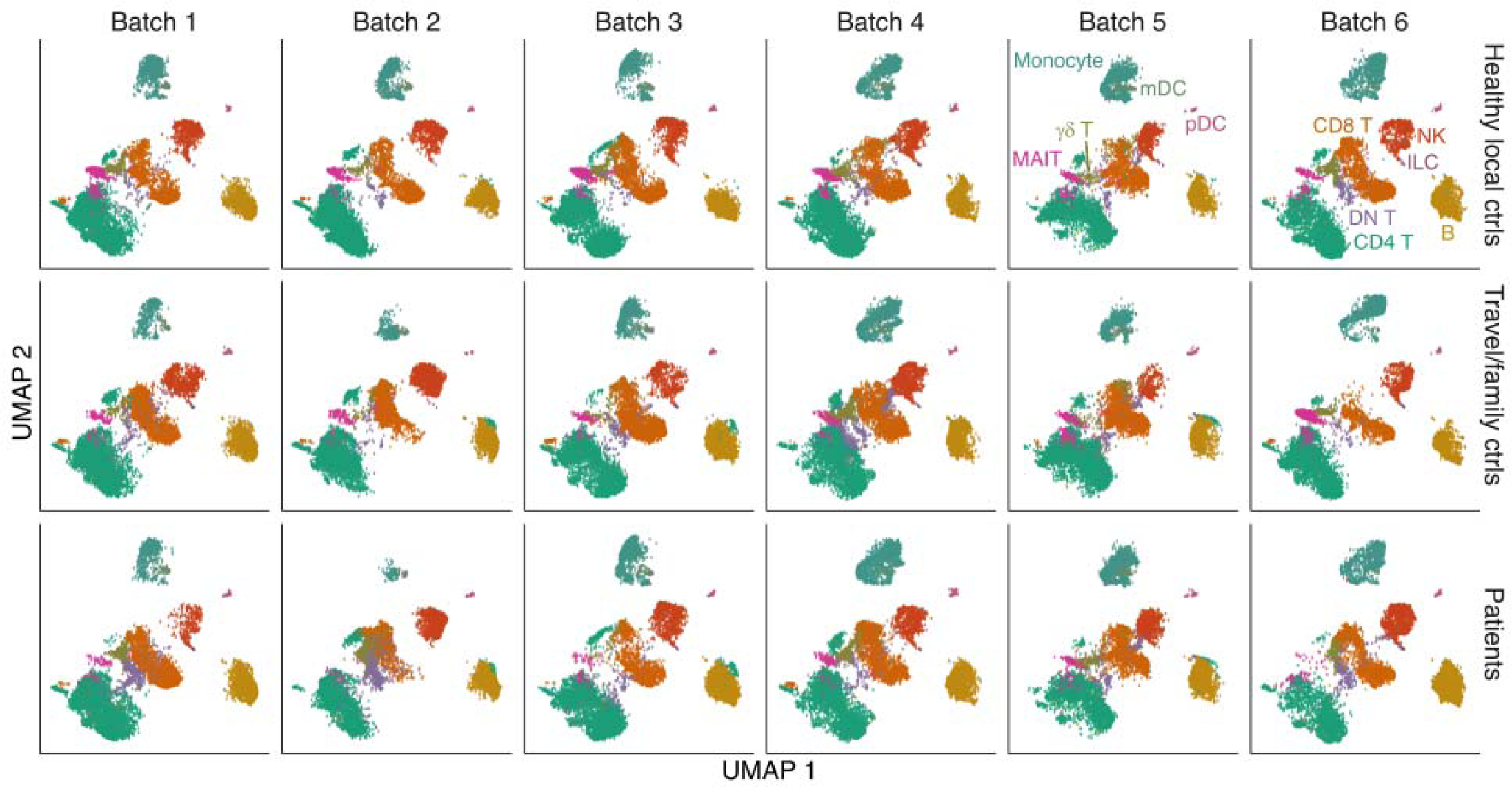
Batch-specific back propagation of cell-type annotations through machine learning. We used 100 cells per cluster to train a batch-specific cell-type classifier by machine learning. Cell types were determined probabilistically for the other cells in a given batch. UMAP dimensions calculated from the non-batch-corrected expression values are shown. The cell types determined by unsupervised clustering followed by manual inspection are indicated by color coding.

**Figure S4.**
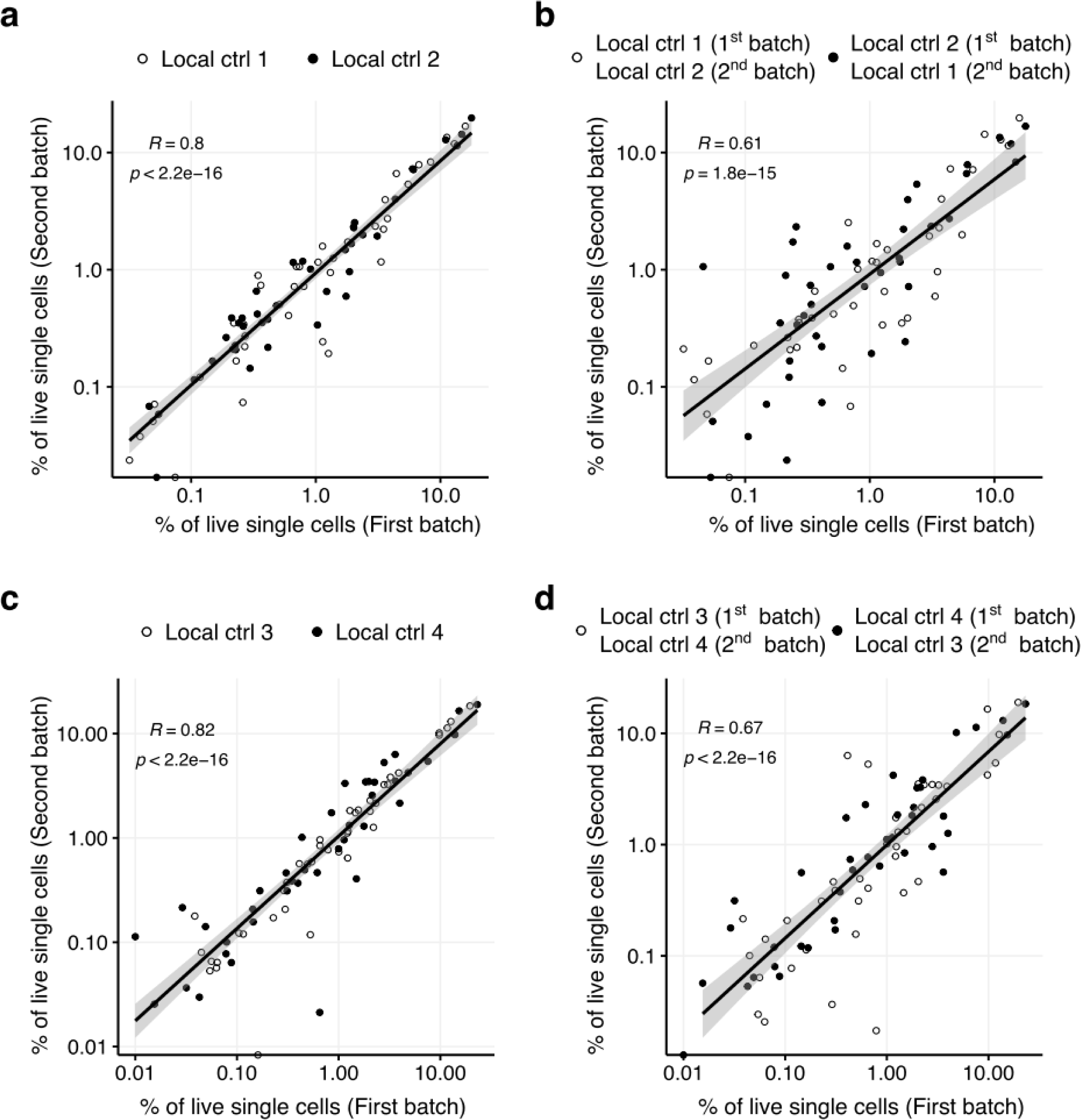
Analysis of the correlation between technical and biological replicates. In our in-house CyTOF datasets, two healthy local controls were analyzed as technical replicates (*i.e*., experiments performed on different dates with aliquots of the same PBMC samples), and another two healthy local controls were analyzed as biological replicates (*i.e*., experiments performed on different dates using PBMC samples obtained on different occasions from the same donors). We assessed the consistency of the predicted immunophenotypes between these replicates, by assessing the correlation between relative frequencies among live single cells (excluding the CD45^-^CD66b^+^ granulocyte-like cluster) of each of the cell types defined by unsupervised clustering followed by manual refinement. We also assessed the “background-level” correlation by intentionally inverting the donors in the second batch. (a) Technical replicates. (b) Dummy analysis of technical replicates in which two donors were intentionally inverted in the second batch. (c) Biological replicates. (d) Dummy analysis of biological replicates in which two donors were intentionally inverted in the second batch. *R*, Kendall’s correlation coefficient.

**Figure S5.**
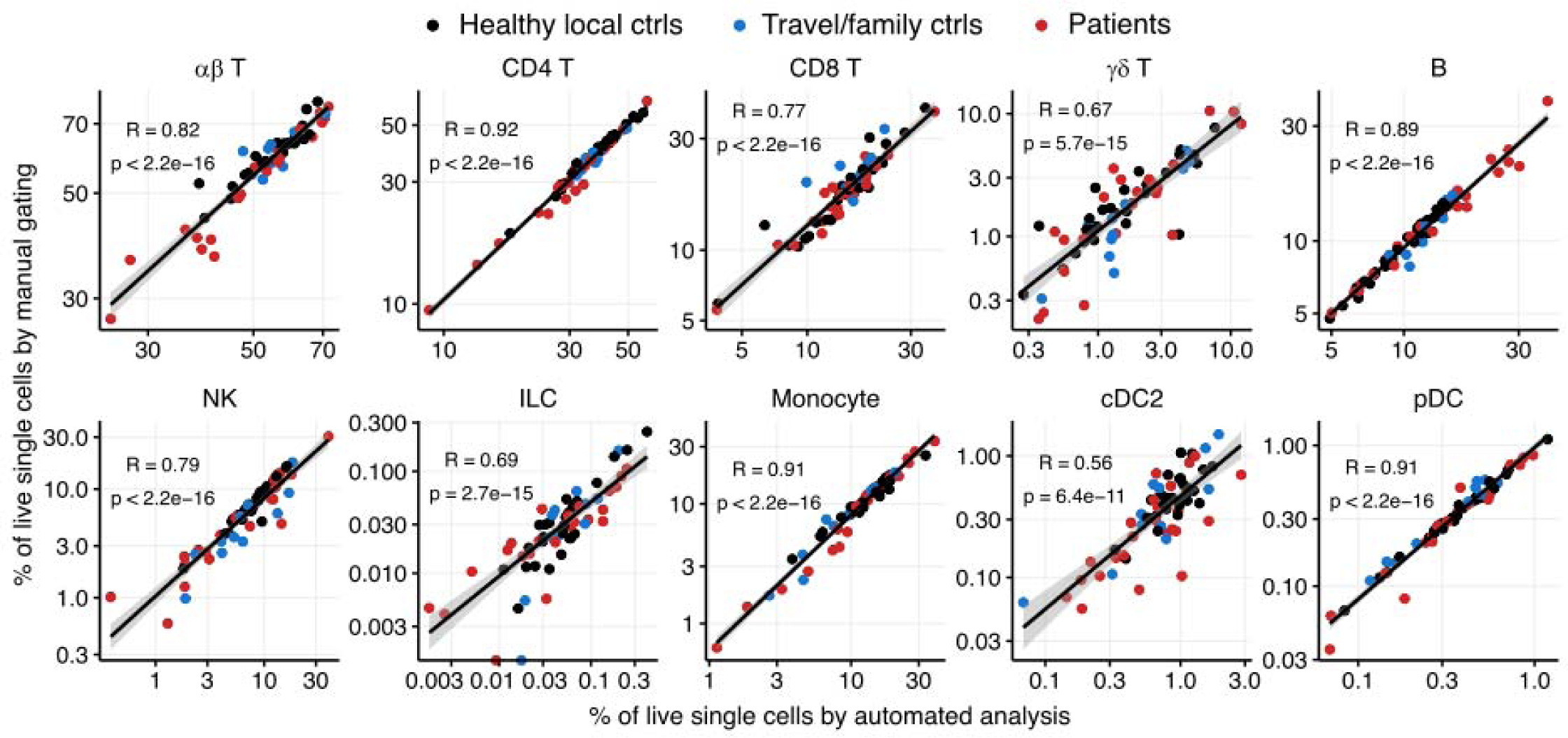
Analysis of the correlation of cell subset abundance values obtained by automated analysis and by manual gating. In-house CyTOF datasets were analyzed with iMUBAC and by manual gating. Frequencies among live single cells were compared for each cell subset. In this analysis, when multiple clusters were identified for a given subset (*e.g*., “NK CD16^+^CD57^-^”, “NK CD16^+^CD57^+^”, “NK CD16^dim^”, and “NK CD56^bright^” for NK cells), these clusters were merged before the calculation of frequencies. Representative results are shown for ten subsets. NK, natural killer cells; ILC, innate lymphoid cells; cDC2, type-2 conventional dendritic cells; pDC, plasmacytoid dendritic cells. *R*, Kendall’s correlation coefficient.

**Figure S6.**
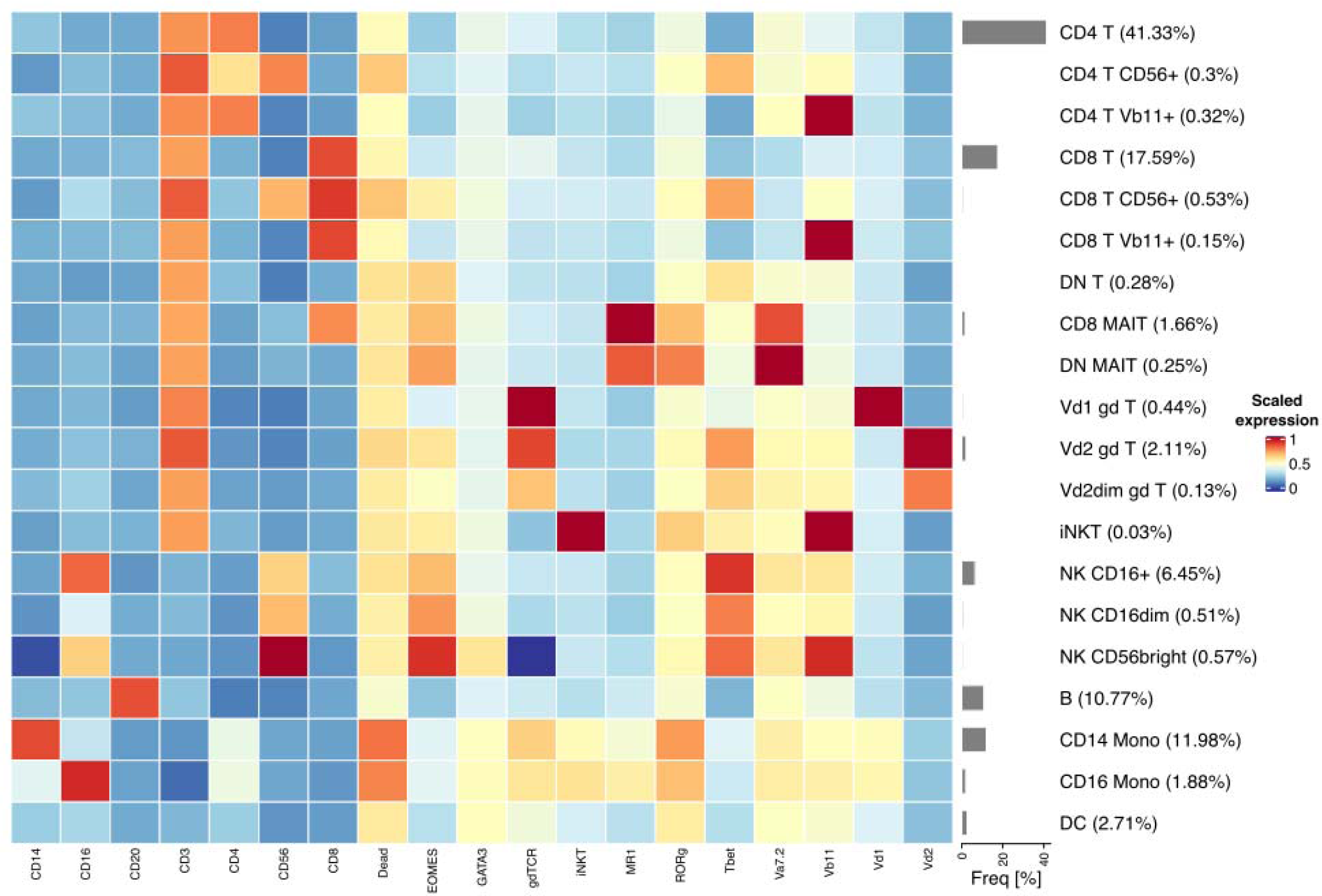
Cell-type identification for the in-house spectral flow cytometry datasets. The datasets consist of data for PBMCs from 11 healthy controls and three patients (two patients with FAS deficiency and one patient with a *STAT3* gain-of-function mutation). The two batches were integrated with Harmony^12^, using default parameters. The FlowSOM/ConsensusClusterPlus method initially identified 60 metaclusters. Clusters were then manually inspected and annotated. A heatmap representing scaled median expression levels is shown.

**Figure S7.**
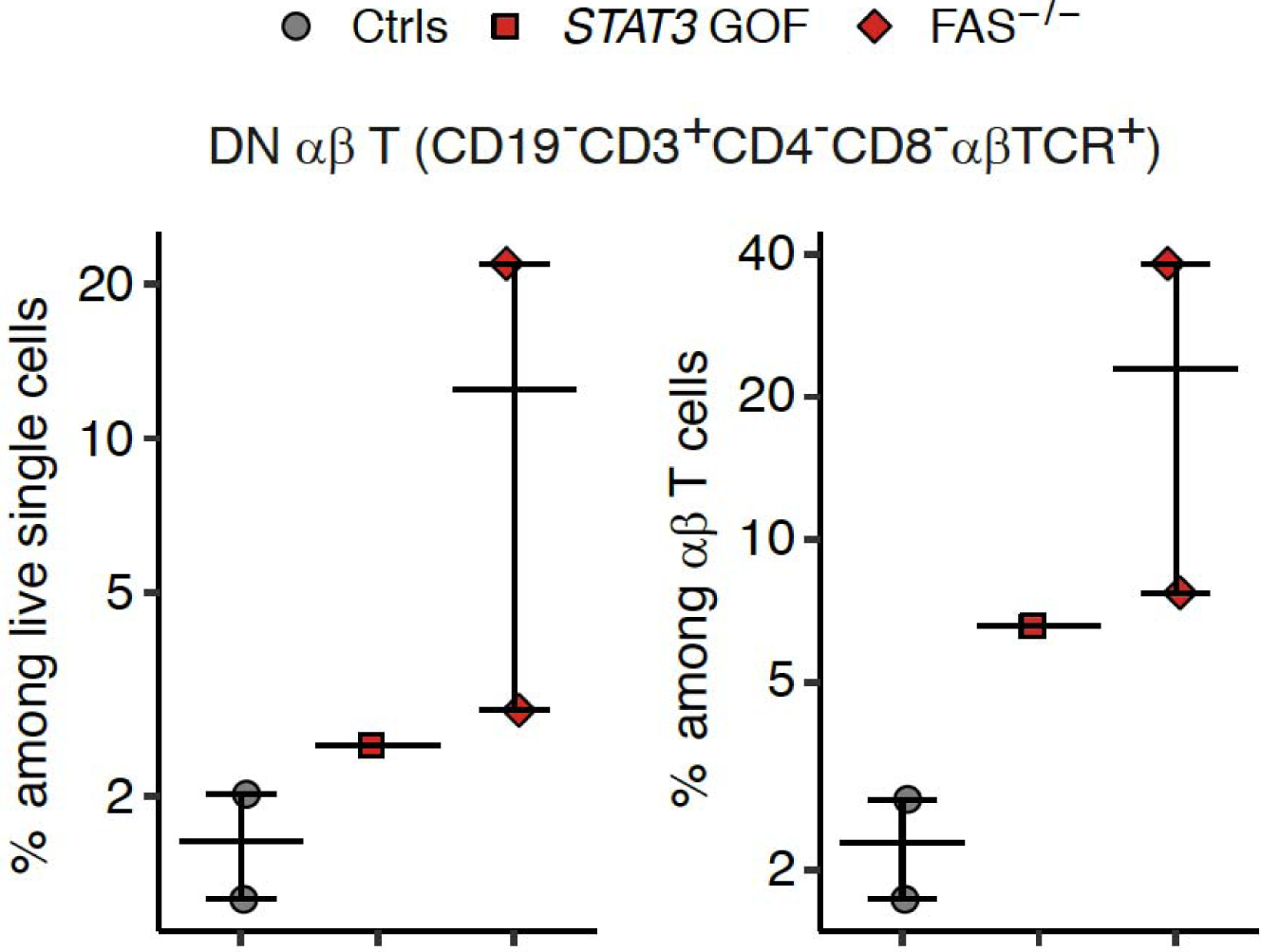
Validation for the in-house spectral flow cytometry data by a conventional flow cytometry. PBMCs from two healthy controls and three patients (two patients with FAS deficiency and one patient with a *STAT3* gain-of-function mutation) were analyzed using a conventional flow cytometry. Compensation was performed using single-stained PBMCs as controls.

**Figure S8.**
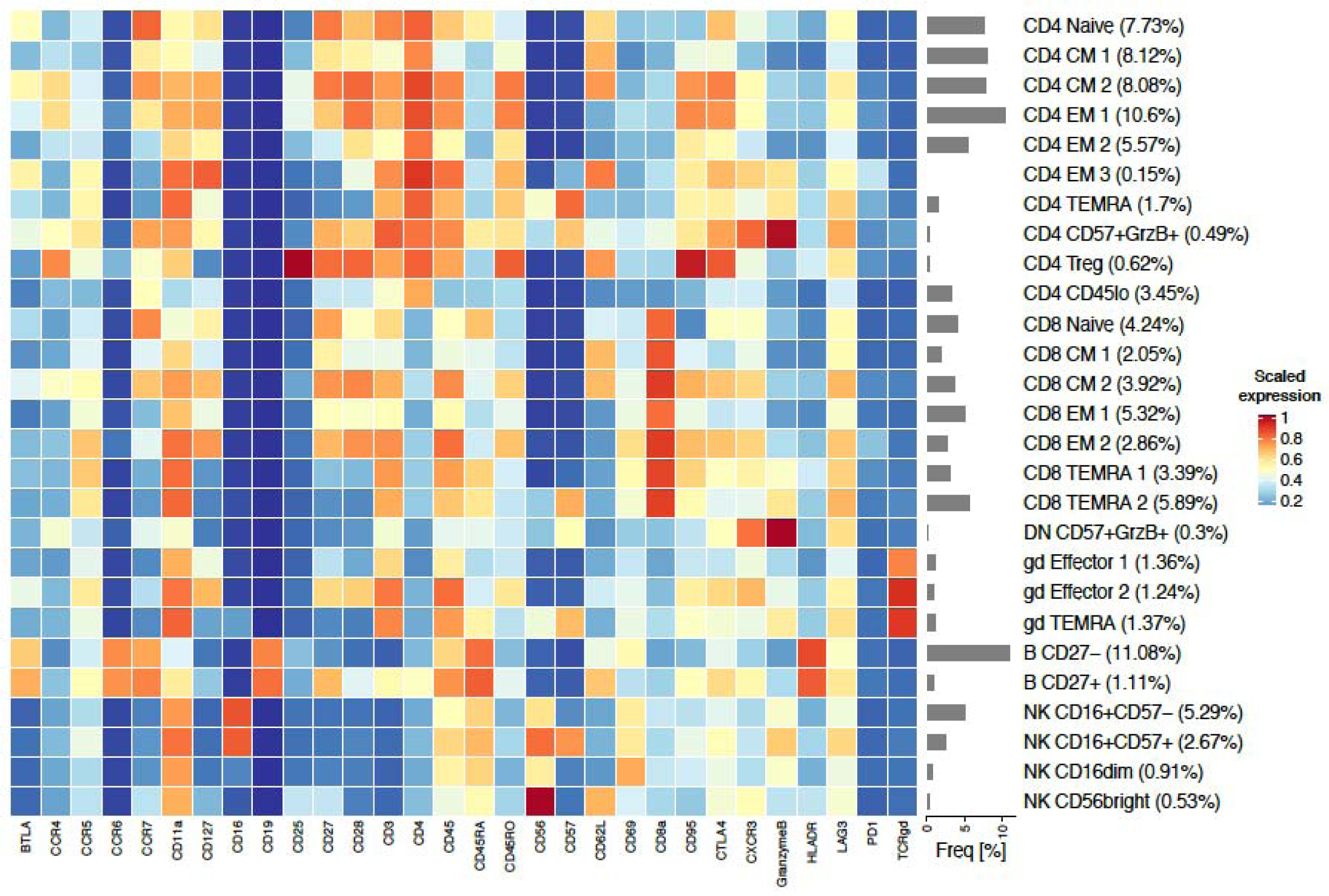
Cell-type identification for the CyTOF Panel 1 dataset for patients treated with PD-1 blockade immunotherapy. The datasets were downloaded from the FlowRepository (FR-FCM-ZY34)^20^. The datasets consist of data for PBMCs from 10 healthy controls and 20 patients with stage IV melanoma, before and after PD-1 blockade immunotherapy (*N*=11 and 9 for responders and non-responders, respectively). The two batches (*i.e*., discovery and validation cohorts) were integrated with Harmony^12^, using default parameters. The FlowSOM/ConsensusClusterPlus method initially identified 40 metaclusters. These clusters were then manually inspected and annotated. A heatmap representing scaled median expression levels is shown.

**Figure S9.**
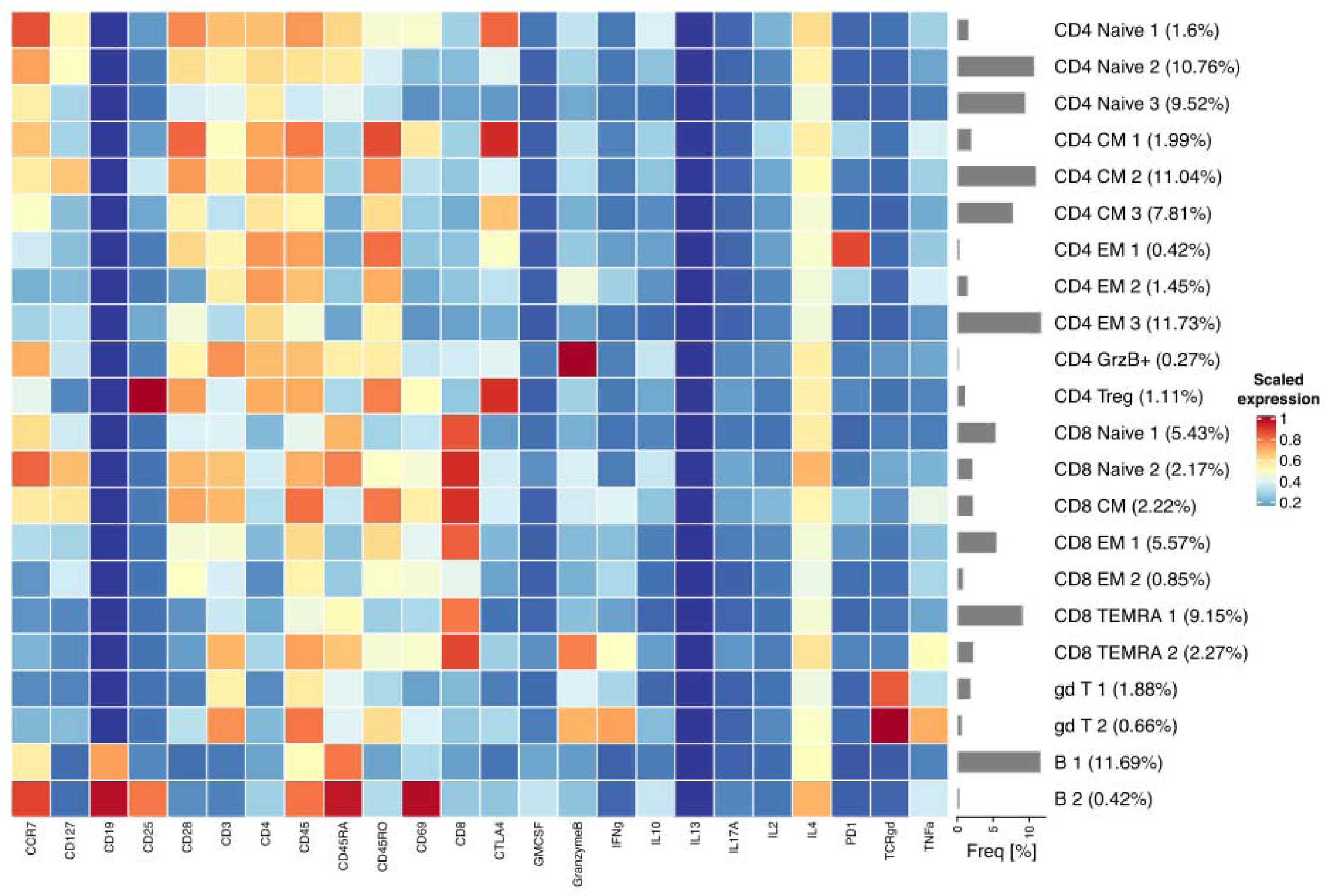
Cell-type identification for the CyTOF Panel 2 dataset for patients treated with PD-1 blockade immunotherapy. See Figure S8 for details.

**Figure S10.**
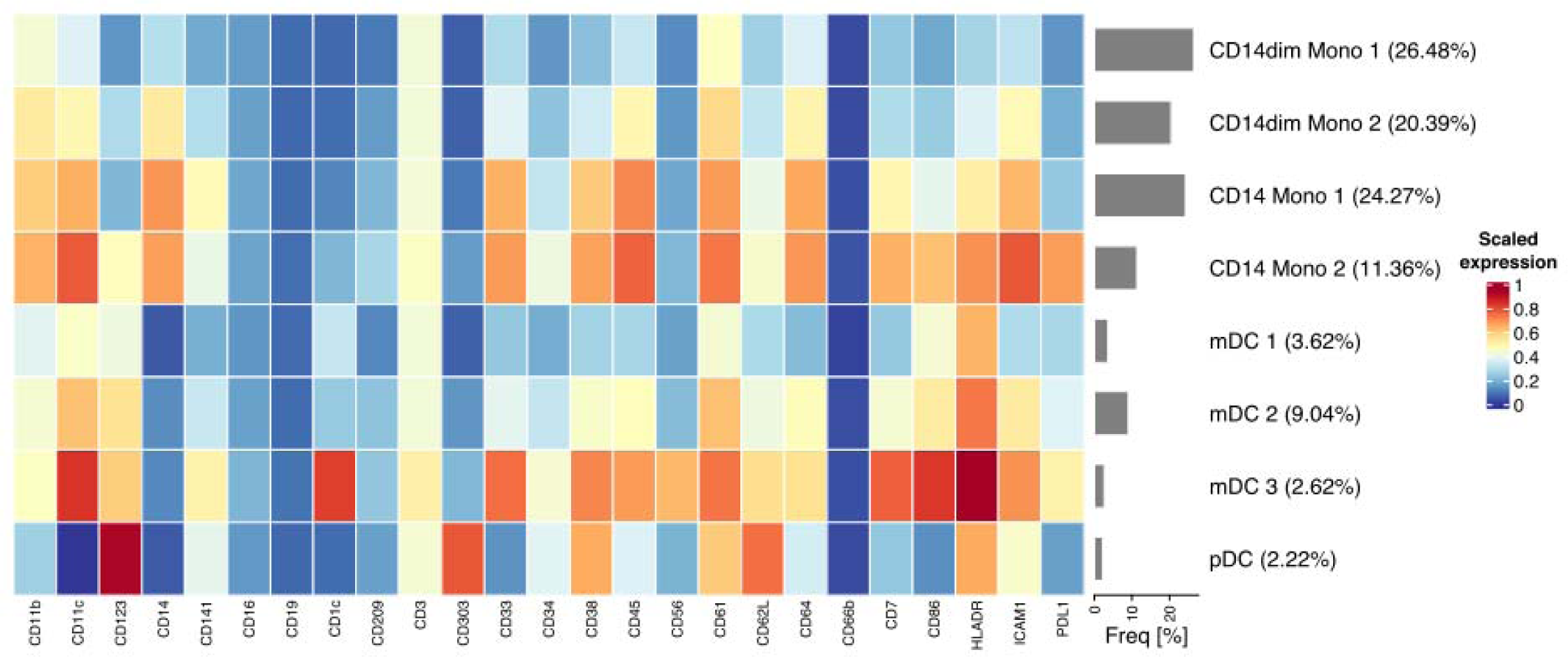
Cell-type identification for the CyTOF Panel 3 dataset for patients treated with PD-1 blockade immunotherapy. See Figure S8 for details.

## Notes

https://github.com/casanova-lab/iMUBAC

### Competing Interest Statement

The authors have declared no competing interest.

## References

1. Tangye, S. G. et al. Human inborn errors of immunity: 2019 update on the classification from the International Union of Immunological Societies Expert Committee. J. Clin. Immunol. 40, 24–64 (2020).

2. Amir, E. D. et al. viSNE enables visualization of high-dimensional single-cell data and reveals phenotypic heterogeneity of leukemia. Nat. Biotechnol. 31, 545–552 (2013).

3. Qiu, P. et al. Extracting a cellular hierarchy from high-dimensional cytometry data with SPADE. Nat. Biotechnol. 29, 886–891 (2011).

4. Van Gassen, S. et al. FlowSOM: Using self-organizing maps for visualization and interpretation of cytometry data. Cytom. Part A 87, 636–645 (2015).

5. Nowicka, M. et al. CyTOF workflow: differential discovery in high-throughput high-dimensional cytometry datasets. F1000Research 6, 748 (2017).

6. Bruggner, R. V., Bodenmiller, B., Dill, D. L., Tibshirani, R. J. & Nolan, G. P. Automated identification of stratifying signatures in cellular subpopulations. Proc. Natl. Acad. Sci. USA 111, E2770–E2777 (2014).

7. Arvaniti, E. & Claassen, M. Sensitive detection of rare disease-associated cell subsets via representation learning. Nat. Commun. 8, 1–10 (2017).

8. Lun, A. T. L., Richard, A. C. & Marioni, J. C. Testing for differential abundance in mass cytometry data. Nat. Methods 14, 707–709 (2017).

9. Van Gassen, S., Gaudilliere, B., Angst, M. S., Saeys, Y. & Aghaeepour, N. CytoNorm: a normalization algorithm for cytometry data. Cytom. Part A 97, 268–278 (2020).

10. Schuyler, R. P. et al. Minimizing batch effects in mass cytometry data. Front. Immunol. 10, 2367 (2019).

11. Amodio, M. et al. Exploring single-cell data with deep multitasking neural networks. Nat. Methods 16, 1139–1145 (2019).

12. Korsunsky, I. et al. Fast, sensitive and accurate integration of single-cell data with Harmony. Nat. Methods 16, 1289–1296 (2019).

13. McInnes, L., Healy, J. & Melville, J. UMAP: Uniform manifold approximation and projection for dimension reduction. 1802.03426. (2018).

14. Xu, C. & Su, Z. Identification of cell types from single-cell transcriptomes using a novel clustering method. Bioinformatics 31, 1974–1980 (2015).

15. Oliveira, J. B. et al. Revised diagnostic criteria and classification for the autoimmune lymphoproliferative syndrome (ALPS): report from the 2009 NIH International Workshop. Blood 116, e35–e40 (2010).

16. Magerus-Chatinet, A. et al. FAS-L, IL-10, and double-negative CD4^-^CD8^-^ TCR α/β^+^ T cells are reliable markers of autoimmune lymphoproliferative syndrome (ALPS) associated with FAS loss of function. Blood 113, 3027–3030 (2009).

17. Haapaniemi, E. M. et al. Autoimmunity, hypogammaglobulinemia, lymphoproliferation, and mycobacterial disease in patients with activating mutations in *STAT3*. Blood 125, 639–648 (2015).

18. Milner, J. D. et al. Early-onset lymphoproliferation and autoimmunity caused by germline *STAT3* gain-of-function mutations. Blood 125, 591–9 (2015).

19. Nabhani, S. et al. *STAT3* gain-of-function mutations associated with autoimmune lymphoproliferative syndrome like disease deregulate lymphocyte apoptosis and can be targeted by BH3 mimetic compounds. Clin. Immunol. 181, 32–42 (2017).

20. Krieg, C. et al. High-dimensional single-cell analysis predicts response to anti-PD-1 immunotherapy. Nat. Med. 24, 144–153 (2018).

21. Robinson, M. D., McCarthy, D. J. & Smyth, G. K. edgeR: A Bioconductor package for differential expression analysis of digital gene expression data. Bioinformatics 26, 139–140 (2009).

22. Weber, L. M., Nowicka, M., Soneson, C. & Robinson, M. D. diffcyt: Differential discovery in high-dimensional cytometry via high-resolution clustering. Commun. Biol. 2, 1–11 (2019).

23. R Core Team. R: A language and environment for statistical computing. (2018).

24. Csárdi, G. & Nepusz, T. The igraph software package for complex network research. InterJournal Complex Sy, 1695 (2006).

25. Kuhn, M. Building predictive models in R using the caret package. J. Stat. Softw. 28, 1–26 (2008).

26. Geurts, P., Ernst, D. & Wehenkel, L. Extremely randomized trees. Mach. Learn. 63, 3–42 (2006).

